# RACLET: the Ramp Above Critical Level Endurance Test to evaluate critical power in cycling

**DOI:** 10.1101/2025.07.11.664347

**Authors:** Maximilien Bowen, Pierre Samozino, Mylène Vonderscher, Baptiste Morel

**Author notes:** This research has been funded by the Agence Nationale de la Recherche [ANR-22-CE14-0073].

## Abstract

This study introduces the Ramp Above Critical Level Endurance Test (RACLET), a novel method for evaluating critical power model parameters, and tests its reliability and concurrent validity. Twenty-three participants completed several RACLET and time-to-exhaustion tests (TTE). The RACLET involves a decreasing power ramp with intermittent maximal sprints, inducing moderate fatigue without exhaustion. The test demonstrated excellent reliability for the initial power (*P*_*i*_) and critical power (*P*_*c*_) (ICC *>* 0.97), with acceptable reliability for the time constant (*τ*) (ICC = 0.70). The concurrent validity against TTE was excellent for *P*_*i*_ and *P*_*c*_ (systematic errors: 1.7% and 3.0%, respectively), but moderate for *τ* and maximum work above the critical power (systematic errors ≈ 10%). The RACLET showed excellent predictive capacity for time-to-exhaustion (systematic error = −0.6%; random error = 10.3%). These results suggest that the RACLET is a reliable, valid, and efficient alternative to traditional critical power testing methods, offering comparable accuracy with a single test without inducing exhaustion. This approach could be particularly beneficial for populations in which exhaustive testing is challenging or impractical.

## Introduction

The concept of Critical Power (P_*c*_) proposes a model that historically describes the relationship between the exercise intensity and the duration or distance that can be sustained. The general form of this decreasing and converging relationship was identified in the early 20th century in racing animals (Kennelly, 1906) and applied to human athletic records (Hill, 1925) before being mathematically formalised half a century later (Monod and Scherrer, 1965). This has since become a cornerstone of exercise physiology (Poole et al., 2016). The Critical Power concept allows for the characterisation of the critical power (*P*_*c*_) as a threshold in biological function (Poole et al., 2016).

Below this threshold, a physiologically stable state can be achieved, but exercising above *P*_*c*_ disrupts homeostasis. A conceptual work reserve (*W ′*) is consumed, and the dramatic onset of fatigue rapidly leads to exhaustion when *W* ^*′*^ is emptied (i.e. the required power is no longer sustainable). The concept of Critical Power has numerous practical applications, including optimising athletes training programs and performances, as well as improving the quality of life of individuals with chronic diseases (Meyler et al., 2025; Muniz and Meyler, 2025; Poole and Jones, 2023).

The historic experimental method, considered the gold standard, consists of performing a series of efforts (usually three to five) at several constant Bowen et al. RACLET – Power power levels and measuring the duration (*t*_*lim*_) before this intensity is no longer sustainable by the participant (Muniz-Pumares et al., 2019). The adjustment of the Critical Power model 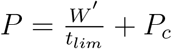 to the data then enables *P*_*c*_ and *W ′*determination. Alternatively, *P*_*c*_ and *W ′* may be estimated using a 3-min all-out test (Vanhatalo et al., 2007), where the mean end power output during the final 30 s of the test represents *P*_*c*_ and the amount of work performed above *P*_*c*_ corresponds to *W* ^*′*^.

However, these conventional methods have major experimental and methodological limitations. Typically, the time-to-exhaustion (TTE) exhibits a high variability of ≈ 15% (Barbosa et al., 2014; Laursen et al., 2007; Muniz-Pumares et al., 2017). This can be attributed to the difficulty in objectively determining the exact time at which the target power is no longer maintained, either because of inherent human variability around the target (Vanhatalo et al., 2007), or psychological aspects (motivation, mental fatigue, familiarisation, Hill, 1993; Salam et al., 2018). Moreover, the fact that these tests lead to exhaustion makes them extremely complicated to implement in fragile populations (e.g. elderly people or pathological populations) or for in-season routine testing in athletes. In the research context, the number of tests required to study the effect of a factor on *P*_*c*_ or *W* ^*′*^ must be multiplied by the number of conditions tested. For example, the effects of heat (Racinais et al., 2015), hypoxia (Townsend et al., 2017), and cadence (Barker et al., 2006) on the P_*c*_ model parameters required 8 to 15 TTE tests. This makes such studies rare, with few participants, likely biased by the high number of sessions, or even impossible to conduct if the number of conditions is excessive.

Richard Hugh Morton has dedicated part of his research activity to mathematically playing with models around the concept of critical power (Morton, 1990; Morton, 1985; Morton, 1986a; Morton, 1986b). Some of these works have become references in the field (Morton, 2006), while others, although containing extremely interesting elements, have almost fallen into oblivion (Morton and Billat, 2013). In this respect, the link between the partial consumption of *W* ^*′*^ and the reduction in maximal capacity (i.e. fatigability) has never been explored. Building on this pioneering work, we recently proposed a mathematical model of exercise fatigability (Bowen et al., 2024). First, this model makes it possible to predict the fatigability (i.e. alteration in maximal capacities) from the known parameters (i.e. *P*_*i*_, *P*_*c*_, and *τ*) for any type of exercise performed in the severe domain. Even more interestingly, this mathematical model also enables the determination of the three parameters by measuring fatigability during exercise.

Taking advantage of this model, it would be possible to design a single test that would induce only moderate fatigue (i.e. without reaching exhaustion) to determine the critical power model’s parameters. A simple idea would be that the participant performs a decreasing ramp starting above and ending below the *a priori* unknown critical power. Although inducing fatigue at the beginning, the power target crosses the critical intensity at some point during the test, allowing the participant to recover. By definition, maximal capacities should decrease during the first fatiguing part of the test and then increase again once the target has crossed *P*_*c*_. Frequent measurements of maximal capacities during the test should therefore make it possible to identify *P*_*c*_ being by definition the target power at which the *fatigue*–*recovery* state transition occurs (Burnley et al., 2012).

Thus, this study aimed to propose a new test, the Ramp Above Critical Level Endurance Test (RACLET), based on the recently developed mathematical model linking fatigability to the Critical Power model. It was hypothesised that this test will enable to reliably and accurately evaluate the parameters of the Critical Power model while inducing only moderate fatigue. Once theoretically developed, a pedaling protocol was set up to test: (i) the RACLET inter-day reliability, (ii) the RACLET concurrent agreement with the gold standard method (i.e. multiple time-to-exhaustion tests), and (iii) the RACLET ability to predict times to exhaustion (i.e. the historical readout of the critical power model).

## Theoretical background

### Model development

Based on the model previously proposed by Bowen et al. (2024), this section aims to model the theoretical change in the maximal pedaling torque capacities as a function of time, Γ_max_(*t*), during an iso-cadence torque-decreasing pedaling ramp exercise. The change in the maximal torque capacity Γ_max_(*t*) during an exercise performed above the critical torque Γ_*c*_ decreases linearly with the integral of the torque above the critical torque with 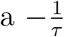 slope. If the initial, free-fatigue torque capacity is Γ_*i*_, the maximal torque capacity at any time Γ_max_(*t*) is defined as:

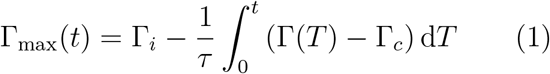

The RACLET is designed such that the intensity starts from a submaximal but supra-critical torque (Γ_*c*_ *<* Γ^⋆^ ≤ Γ_*i*_) and decreases linearly with slope *S* (Eq. 2) :

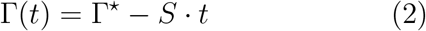

By combining Eq. 1 and Eq. 2, the maximal torque capacity Γ_max_(*t*) evolves as a quadratic function of time:

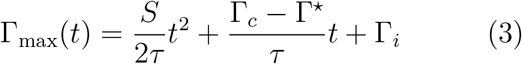

Thus, by developing the generic model (Eq. 1) for a decreasing ramp exercise, the theoretical decrease in maximum torque capacity Γ_max_(*t*) is described, as long as Γ(*t*) ≥ Γ_*c*_, by a second-order polynomial function, depending on three parameters: the initial torque Γ_*i*_, the critical torque Γ_*c*_, typical time constant *τ* and the two parameters of the ramp exercise (Γ^⋆^ and *S*). *Nota bene*, in the specific case of an iso-cadence exercise, the crank angular velocity (*ω*) is constant, so power *P* = *ω* ·Γ is equivalent to Γ (*P* ≡ Γ) and Eq. 1, Eq. 2 and Eq. 3 can be expressed, by analogy, as:

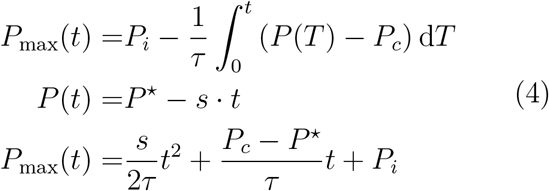

with the initial power *P*_*i*_, critical power *P*_*c*_, the power ramp start *P* ^⋆^, and slope *s* being equivalent to Γ_*i*_, Γ_*c*_, Γ^⋆^ and *S* in terms of power. Thus, as long as *P* (*t*) ≥ *P*_*c*_, the maximal *P*_max_(*t*) can be expressed as a 2nd order polynomial function *P*_max_(*t*) = *A · t*^2^ + *B · t* + *C*, where the three parameters are:

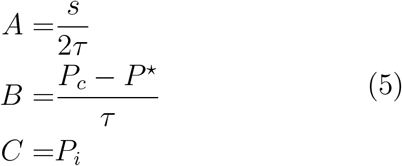

Based on this model, if a participant follows the linearly decreasing target power *P* (*t*) (Eq. 4) and regularly performs maximum pedal strokes, his maximum power is assumed to start from *P*_*i*_ and then decrease, until the target power *P* = *P*_*c*_. At this specific time point (*t*_*c*_), the exercise is no longer assumed to generate fatigue, mathematically,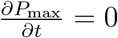. Immediately thereafter, *P* (*t*) *< P*_*c*_, and recovery (i.e. *P*_max_ increases) should be observed (Morton, 2006), although mathematically outside the applicability of the model constrained to the severe domain. The theoretical evolution of the maximal power *P*_max_ during the decreasing ramp *P* (*t*) and a graphical representation of the parameters are displayed in Fig. 1.

**Figure 1.**
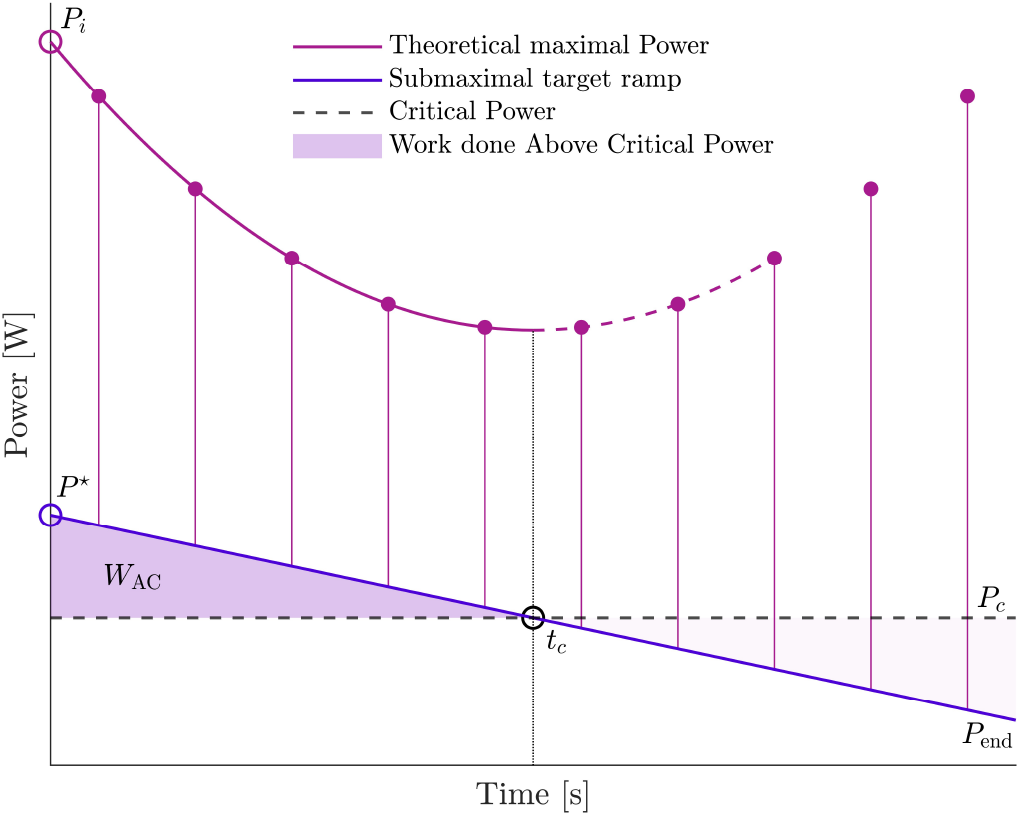
Theoretical representation of the Ramp Above Critical Level Endurance Test (RACLET). The blue line represents the target power starting at *P*^⋆^ and ending at *P*_*end*_. The maximum power capacity was measured regularly using maximal pedal strokes (purple dots). During the first phase of the test, a work above the critical power is produced (*W*_AC_; shaded area). At this point, the participant is in the fatigue phase, and the maximum power, starting at *P*_*i*_, decreases according to a theoretically quadratic law (purple line; Eq. 4). At an a priori unknown time *t*_*c*_, the target power will cross the critical power *P*_*c*_, and the participant will then start the recovery phase; the maximum power capacity will increase accordingly. Note that the model is displayed as a dotted purple line as it was not built to describe the recovery phase.

### RACLET settings

To operate experimentally, the parameters of the RACLET ramp exercise (Γ^⋆^ and *S*) must be individually determined to meet a series of constraints: (i) the RACLET must initially fatigue to generate a decrease and then a recovery in *P*_max_; the local minimum of the *P*_max_(*t*) relationship is the key to determining *P*_*c*_; (ii) the RACLET must not generate excessive fatigue (*P*_max_(*t*) always above *P* (*t*)) to avoid exhaustion and the inability to complete the test; and (iii) maximum capacity assessments must be frequent enough to adjust the model, but not too frequent to limit its impact on fatigue. Based on these constraints, we propose below a framework to determine the RACLET settings, in particular the power ramp start *P* ^⋆^ and slope *s*.

*W*_AC_, corresponding to the accumulated work above the critical power, is directly related to fatigue and is mathematically defined as the integral of *P* (*t*) *> P*_*c*_. Thus, during a RACLET:

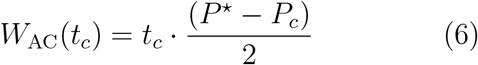

where *t*_*c*_ denotes the instant at which *P* reaches *P*_*c*_ (i.e. *P* (*t*_*c*_) = *P*_*c*_). The parameter *t*_*c*_ can be expressed as a percentage *γ* of the total duration of the RACLET (*t*_*R*_), as: *t*_*c*_ = *γ* ·*t*_*R*_, with 0 ≤*γ* ≤ 1. Combined with Eq. 6, *P* ^⋆^ can be expressed as a function of *W*_AC_:

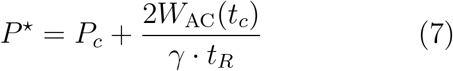

To set the target fatigue level of the RACLET, *W*_AC_(*t*_*c*_) can be defined as 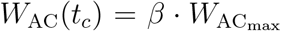, where *β* is the fraction of 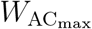 consumed during the test at *t* = *tc*; thus, the relative fatigue level is restricted to 0 ≤ *β* ≤ 1. *P*_*c*_ can also be expressed as a ratio of *P*_*i*_; thus, *P*_*c*_ = *α· P*_*i*_ where 0 ≤ *α* ≤ 1. Similarly, the initial power target *P* ^⋆^ can be expressed relatively to the maximum capacity as: 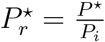 with 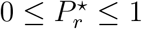.

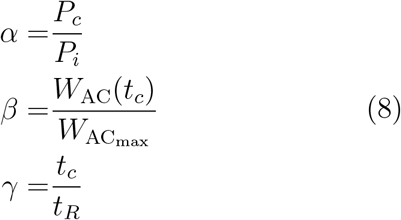

So, the relative initial power of the ramp 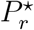 can be experessed as:

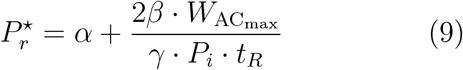

And, the relative ramp slope *S*_*r*_ as:

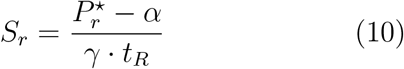

From Eq. 7, it is possible to set the initial intensity of the RACLET *P* ^⋆^ by fixing *t*_*R*_ (the total duration of the test), *γ* (the ratio between *t*_*c*_ and *t*_*R*_, i.e. the duration of the fatiguing part of the test relative to the total duration), *β* (the fatigue level targeted at *t* = *t*_*c*_), and with the expected characteristics of a participant or population being (i) the maximal power *P*_*i*_; (ii) *α*, the critical power expressed relatively to *P*_*i*_, and 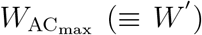, the maximal work capacity above the critical power *P*_*c*_. These data can be obtained from pilot testing or previously published data. Note that *γ* and *β* are *a priori* values that will probably differ experimentally in regard to the actual characteristics of the participants.

## Methods

### Participants and experimental design

Twenty-three volunteers participated in this study (8 females and 15 males; mean±s.d.; age: 23 ± 4 years; mass: 69.2 ±10.7 kg; height: 175.1 ± 10.2 cm). The experiment was conducted over five sessions 24 to 48 h apart in order to test the pedaling task (i) the RACLET reliability (sessions 1 and 2 conducted identically; hereafter referred to as RACLET (A) and (B), and (ii) the RACLET concurrent agreement with the multiple time-to-exhaustion gold standard method (sessions 3 to 5). All participants were healthy and physically active, without injuries and without any drug, medication, or nutritional supplement consumption that could have influenced the test. Written informed consent was obtained from all participants and the study was conducted in accordance with the Declaration of Helsinki. Approval for this project was obtained from the local Ethics Committee on Human Experimentation.

### Material (*Apparatus*)

All tests were performed using a customised iso-inertial cycle ergometer (Monark LC6, Stockholm, Sweden) instrumented with a strain gauge with its amplifier (Futek FSH04207; 444.822

N; M6×1; 500Hz Gain 1.9N/V) that measures the frictional force applied by the belt to the flywheel, and with an optical encoder (Hengstler 600 points per revolution) that assesses the angular displacement of the flywheel. The friction force applied to the belt was controlled using a motorised linear actuator. The external torque (in N · m) and cadence (in rpm) produced by the participant at the crank were calculated using the gear ratio (52/14), radius of the wheel (0.257 m), and torque to overcome the flywheel inertia (I=1.01 kg ·m2, Arsac et al., 1996). Torque and cadence signals were sampled (200 Hz) and stored in a custom LabVIEW program (National Instruments Corporation, Austin, TX, USA) via a 16-bit analogue-to-digital interface card Ni DAQ (NI-USB6210 National Instruments Corporation).

To ensure that the torque targets were respected during the different tests, the cyclo-ergometer (Fig. 2A.) was equipped with a linear actuator connected to the friction belt surrounding the flywheel. This actuator was driven by a custom PID (Proportional – Integral - Derivative) regulator to adjust the friction force to a target value. The actuator displacement motor is driven by the PID function of the error (*e*) between the desired target torque (*T*) and the measured torque produced (Γ) in Eq. 11 and Eq. 12 (Fig. 2B.).

**Figure 2.**
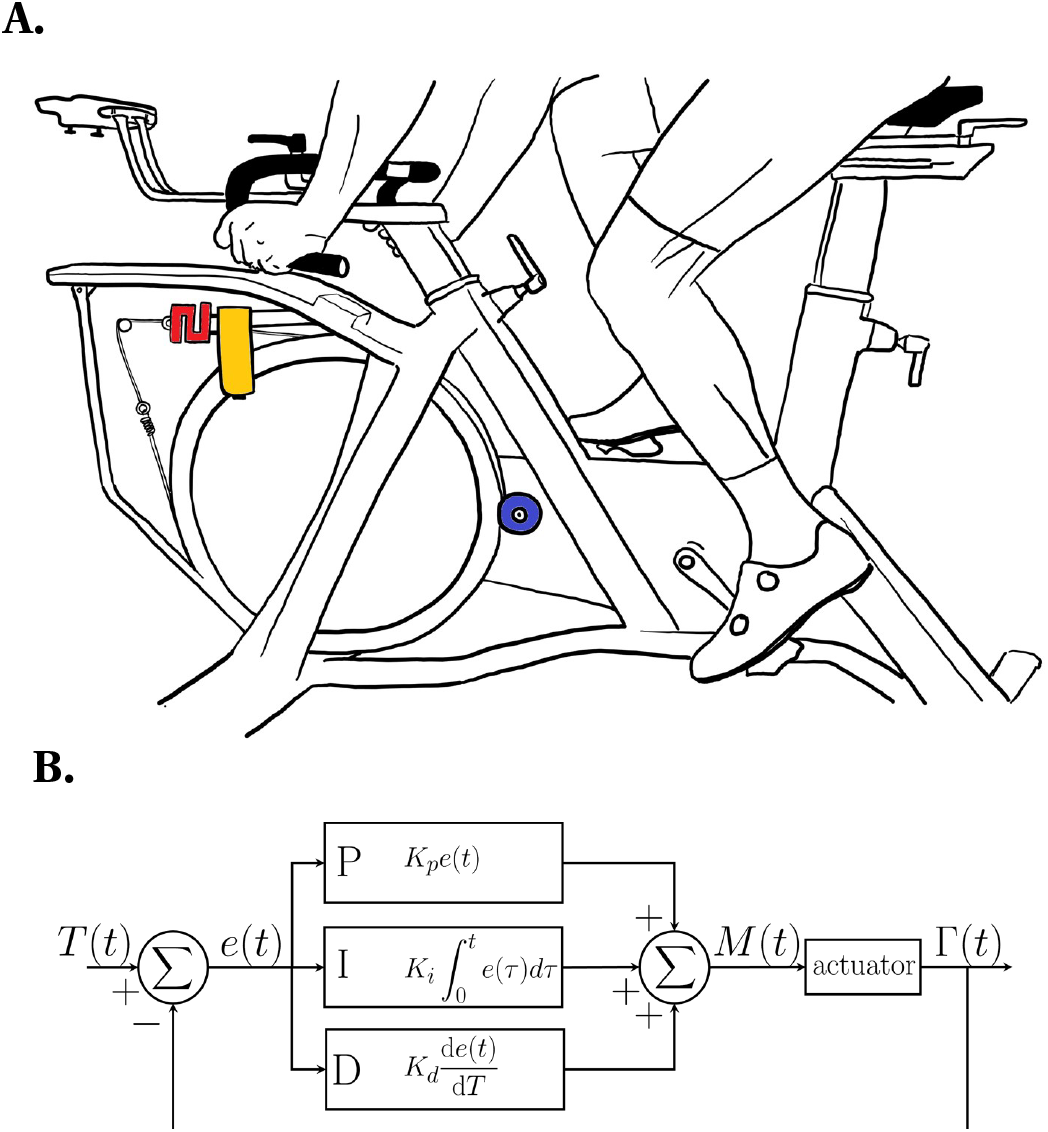
Customised cyclo-ergometer. (**A**) red: strain gauge measuring belt friction force; blue: optical encoder measuring angular velocity of the flywheel; yellow: motor controlling belt friction. (**B**) PID Regulator: The proportional (*K*_*p*_), integrative (*K*_*i*_), and derivative (*K*_*d*_) constants were set as 8.1 · 10^*−*4^, 1.0 ·10^*−*5^, and 1.0 · 10^*−*6^, respectively.

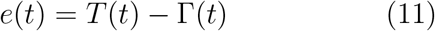

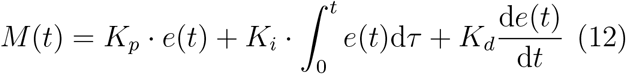

### Protocol

The participants used toe clip pedals and sat on the bike saddle at all times. All tests were performed in an iso-cadence condition requiring the participant to maintain a 80 − rpm cadence as stable as possible, owing to visual feedback (cadence was displayed on a screen) and auditive feedback (metronome set to the pedaling frequency). All efforts requiring maximal intensity (hereafter referred to as sprints) were supported by strong verbal encouragements provided by the investigators. Each experimental session started with a 10 min standardised warm-up, with 5 min of cycling at 1 W*·*kg*−*1 and 5 min at 2 W*·*kg*−*1.

#### RACLET

##### RACLET setting

The test duration *t*_*R*_ was set to 300 s. Ten evaluations of the maximal capacity were deemed necessary to fit the model accurately. Furthermore, limiting the duration of sprints to 10% of the test duration seemed to be a limit that provided acceptable results during the pilot phase (3 s sprint every 30 s for 300 s). Furthermore, the first sprint was performed 10 s after the RACLET start to avoid beginning the test with a maximum effort. The relative fatigue parameter was set to generate a relative level of fatigue at ∼ 50% exhaustion (*β* = 0.5, i.e. a consumption of 50% of 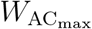) to induce fatigue without leading to exhaustion. Typical values of a healthy, physically active population can be retrieved from the literature: *P*_*i*_= 1130 W (Dorel et al., 2010), *α* = 0.25 (Vanhatalo et al., 2007), and 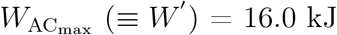 (Vanhatalo et al., 2007). Substituting these parameters into Eq. 9 yields a 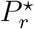 of 0.38.

##### RACLET procedure

First, participants completed three accelerated sprints with friction decreasing in the ramp from 1 to 0 N · kg*−*1 in 6 s (Rozier-Delgado et al., 2025), and interspersed by a 5-min passive recovery. The torque/power-cadence relationship was fitted to these data. The sprint with the best *r*^2^ was retained to determine the maximum torque for the

80 − rpm condition (Γ_80_; Fig. 3). This allows the computation of RACLET’s initial friction torque as Γ^⋆^ = 0.38 *·* Γ_80_.

**Figure 3.**
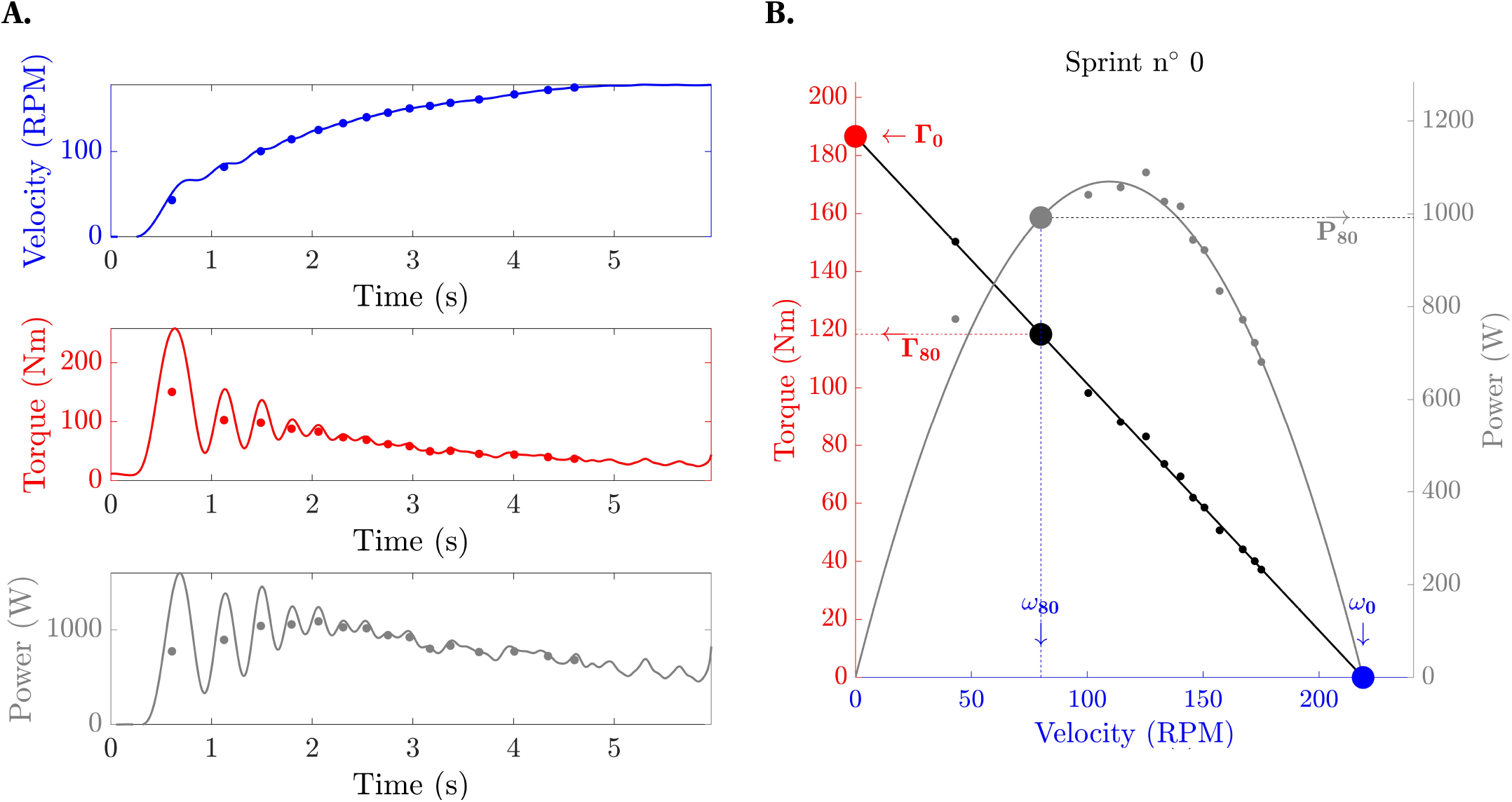
Typical traces of mechanical data and initial Force-Velocity (black) and Power-Velocity (gray) relationships: (**A**) The average value of each pedal stroke in velocity, torque and power (dots) were used to fit Eq. 13. (**B**) The initial maximum torque and power at 80 rpm (Γ_80_ and *P*_80_) weer computed from this relationship.

Following a 5-min recovery period, the participants performed a familiarisation bout to become accustomed to the RACLET testing procedure. This corresponds to the first 100s of the RACLET (including four sprints) with a 5% lower Γ^⋆^ to limit a fatigue effect. After another 5-min of recovery, the participants performed a RACLET protocol consisting of a torque ramp decreasing from Γ^⋆^ to very low intensity (5% of *P*_*i*_) in 300 s; thus, the ramp slope was 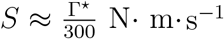. Ten seconds after the start and every 30 s thereafter (total = 10 times), participants were asked to perform a very short sprint of six pedal strokes (i.e. three crank revolutions) performed from ≈60 to ≈100 rpm (total duration ≈3 s) in order to compute the maximum torque and power at 80 − rpm. To achieve this, the friction was briefly increased to brake the flywheel. The participant back-pedaled in the optimal crank position and, without recovery, started sprinting against a decreasing friction (adapted from Rozier-Delgado et al. (2025) with a friction ramp from +150% to 5% of Γ_80_ in 3s). The back pedal was realised to prevent the participant from getting jammed in a dead centre position. After the sprints, the friction instantly returned to the ramp target value. The rate of perceived exertion Bowen et al. RACLET – Power (RPE 6-20 scale) was asked immediately after each sprint, the highest and end-test values were conserved.

#### Time to exhaustion

To test the concurrent agreement between the RACLET and the multiple time-to-exhaustion considered the gold standard method, sessions 3 to 5 consisted of constant power exercises until exhaustion. Three randomised constant powers were calculated for each participant to generate a time-to-exhaustion of approximately 1, 5, and 12 min. These tests were performed with a constant 80-rpm cadence and stopped when the participant could no longer maintain cadence (≤ 75 rpm for 5 consecutive seconds). The mean power and time-to-exhaustion (*t*_lim_) were recorded for further analysis.

### Data analysis

All raw signals were filtered using a low-pass null-phase Butterworth 3rd-order filter at a cutoff frequency of 5 Hz (which removes all non-pedaling high-frequency phenomena, i.e. bike vibrations and electromagnetic noise) and preserves the pedaling signal lower than 300 rpm ≡ 5 Hz, *ceteris paribus*). Torque, power, and cadence were averaged over each stroke, and the timestamps per pedal stroke were recorded.

#### Maximal capacities evaluation

The maximal power capacity at 80 rpm 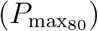, is the maximal capacity regularly evaluated during the RACLET. To do so, a torque-cadence relationship was fitted on each sprint six maximal pedal strokes intended to be centred at 80 rpm (≈ from 60 to 100 rpm, *vide supra*). For each sprint, a pedal stroke was conserved if it presented the highest cadence among pedal strokes with a lower torque and *vice versa*. Subsequently, the Γ(*ω*) function (Eq. 13) was fitted with the remaining torque and cadence data to obtain Γ_0_ and *ω*_0_. 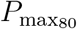 was computed from Eq. 14 (Fig. 4) and conserved in the analysis only if the *r*^2^ of this relationship was *>* 0.5, otherwise, the sprint was discarded.

**Figure 4.**
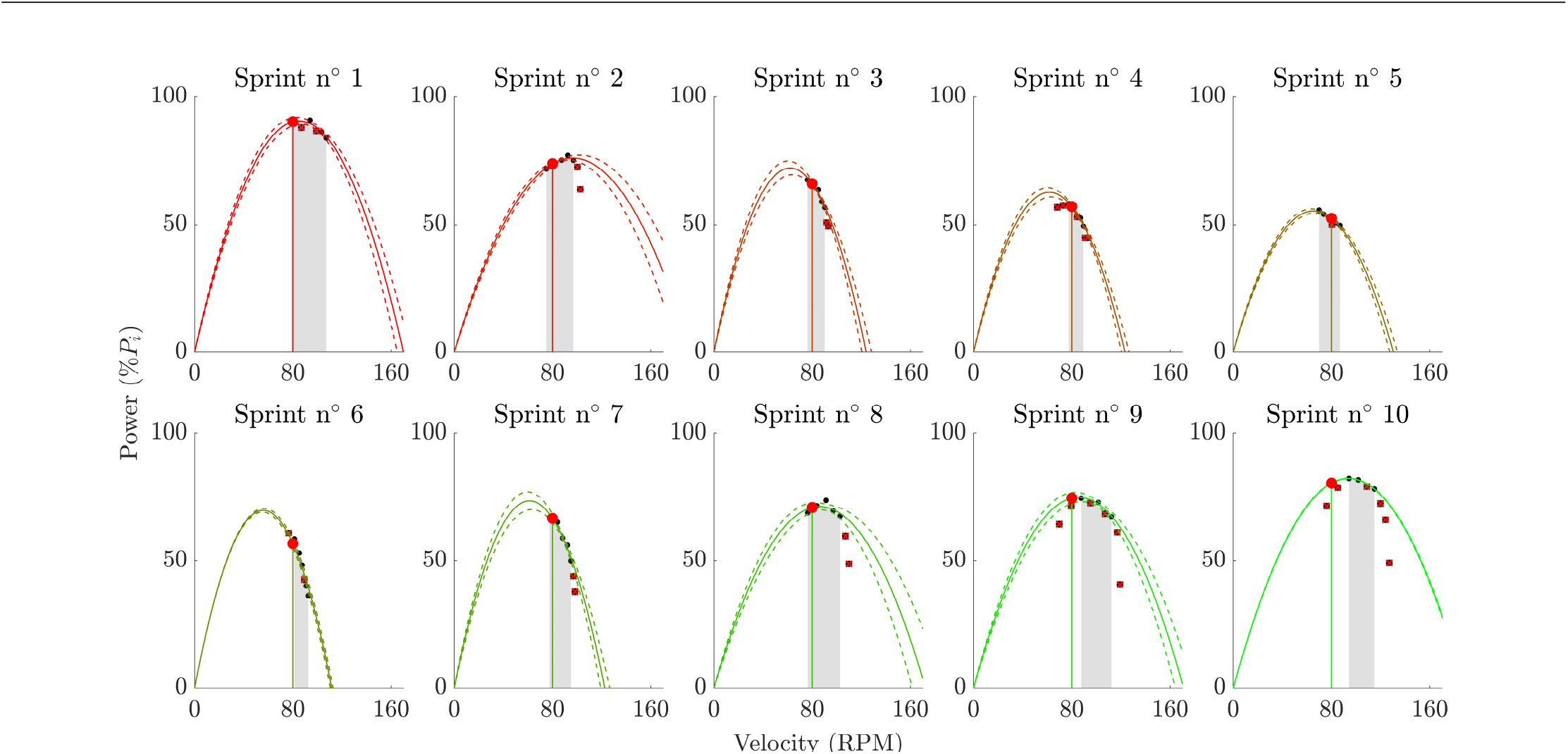
RACLET’s quick sprints Power-Velocity of maximal pedal strokes. Each pedal strokes are displayed with a black dot. Red cross identify outliers removed from the analysis. Red dots represent the maximal power 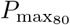 computed by interpolation at 80 rpm (shaded area represents the range of cadences covered during the quick sprint). The colour of the power-velocity relationships indicates the number of sprints (1 to 10) and the state of fatigue (red) or recovery (green).

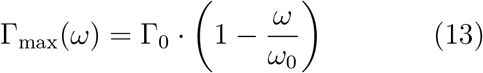

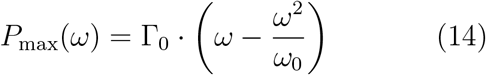

#### Fitting parameters from RACLET

The initial 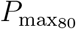 (Sprint n*?*0; Fig. 3) as well as all 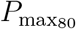 obtained from each sprint during the RACLET (Sprint n*?*1 to 10, Fig. 4) were gathered and associated to a timestamp. These 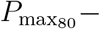 time data were filtered using a Butterworth low-pass filter at 1/150 Hz, which is twice as high as the expected phenomenon (a fatigue-recovery cycle in ≈ 300 s, during RACLET). Only data points up to three after the minimum 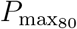 were conserved (Fig. 5A.). As a reminder, the model used was not intended to describe the recovery phase. Three points after the minimum is a trade-off that facilitates the identification of 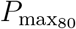 local minimum, i.e. when fatigue stops, allowing recovery, without generating major errors. The actual initial power *P* ^⋆^ and ramp slope *s* were determined by fitting the experimental ramp data (excluding short sprints). Then, Eq. 4 was firstly fitted to the 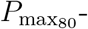 time data using the previously determined *P* ^⋆^ and *s* and a Trust-Region algorithm. Outliers were identified as z-score residuals outside of a ±3 boundary and discarded before a second fitting procedure, which allowed for the estimation of the three parameters *P*_*i*_, *P*_*c*_, and *τ* computed from the RACLET data. Using these three parameters, 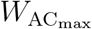 was also computed (Eq. 16). Graphically, the critical power *P*_*c*_ corresponds to the value of the ramp *P* (*t*) at *t*_*c*_ when the minimum theoretical 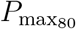 is reached as shown in Fig. 5A., under the iso-cadence condition (Fig. 6B.).

**Figure 5.**
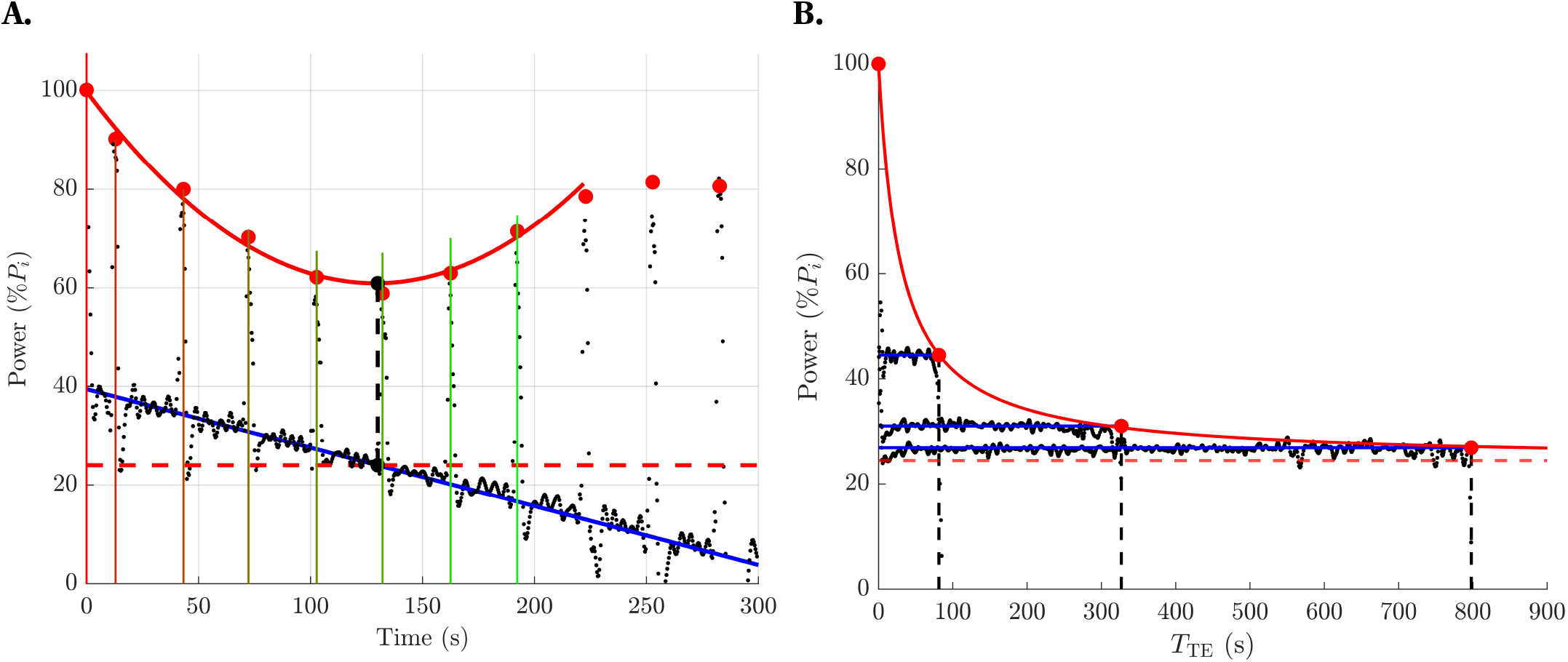
Typical Result obtained for a participant for: (**A**) Power-Time relationship during RACLET at 80 rpm. Small black dots indicate pedal strokes. Red dots: maximal power at 80 rpm 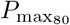. Black dot: local minimum of the maximal power - time relationship. The target power (blue line) at the moment when the maximum power began to recover (red line apex) corresponded to the critical power (dashed red line). (**B**) Power-Duration relationship obtained at 80 rpm. Blue line: target power; Black line; actual power; Red dot: time-to-exhaustion, i.e. duration; Dashed red line : critical power determined from Eq. 15.

**Figure 6.**
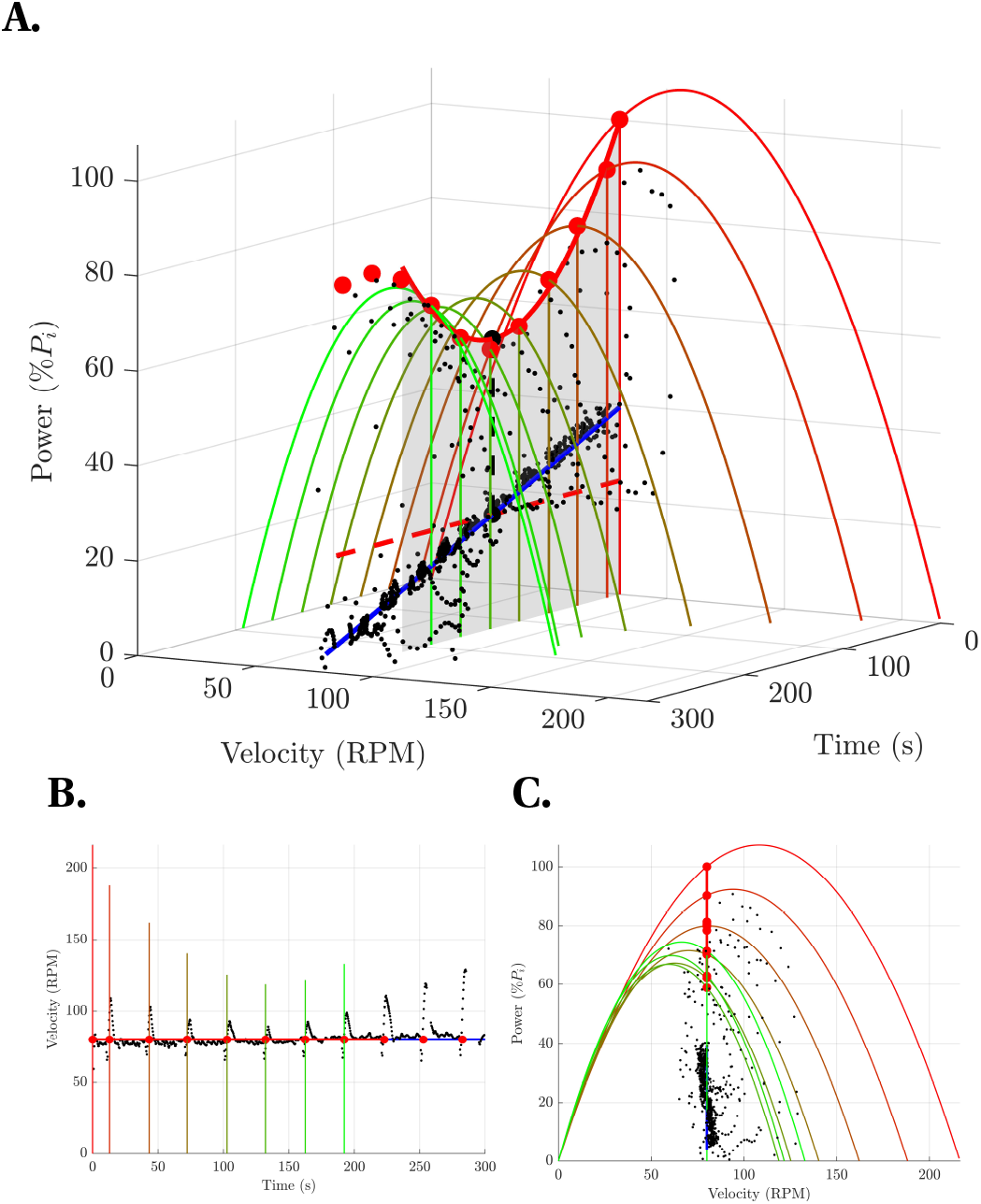
Iso-cadence 80 rpm RACLET typical data represented in 3D Power-Velocity-Time (A) as well as 2D projections of Velocity-time (B) and Power-Velocity (C). Small black dots: pedal strokes. Red dots: maximal power at 80 rpm 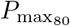. The target power (blue line) at the moment when maximum power begins to recover (red line apex) corresponds to the critical power (dashed red line). The colour of the power-velocity relationships indicates the number of sprints (0 to 10) and the state of fatigue (red) or recovery (green).

#### Fitting parameters from TTE

The three time-to-exhaustion (∼1, ∼5, and ∼12 min) and the 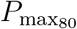 corresponding to the initial power *P*_*i*_ at *t* = 0 were fitted with a 3-parameter model Eq. 15. The three parameters *P*_*i*_, *P*_*c*_, and *τ* were determined from the time-to-exhaustion data (*t*_lim_) and 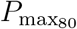, determined by the first sprint.

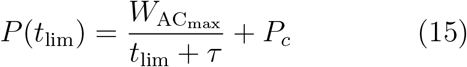

where:

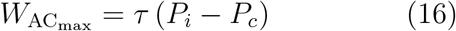

#### Statistical analysis

The data were analysed using a custom-written code in MATLAB Software (2023a) and are presented as mean ± standard deviation. To assess the relative and absolute reliability of the two RACLET, the Intraclass Correlation Coefficient (ICC2,1; Koo and Li, 2016), Typical Error of Measurement (TEM), and Change in the Mean (CM), as well as their respective 95% confidence limits to determine the precision of the estimates, were computed for *P* ^⋆^, *s, P*_*i*_, *P*_*c*_, *τ* and 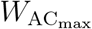. ICC and TEM were considered acceptable if they were greater than 0.7 and less than 10%, respectively (Atkinson and Nevill, 1998). The coefficient of correlation *r* and p − value were also used to calculate the correlation between the two tests.

To test the level of concurrent agreement with the proposed computational method, we compared the parameters obtained from the RACLET versus the gold standard TTE methods. The agreement between the two methods was assessed using the mean difference (i.e. systematic error; SE), and standard deviation of the differences (i.e. random error; RE), which were expressed with 95% confidence limits to determine the precision of the estimates. The coefficient of correlation *r* and p − value were used to calculate the relationships between the critical power values of the two tests. The level of statistical significance was set at p*<*0.05.

Finally, the predictive capacity of the proposed model was tested by comparing the predicted and observed time-to-exhaustion values from the three TTE. Using the model of Bowen et al., 2024, the time-to-exhaustion can be predicted for a given level of work produced *W* = *P* · *t*_lim_ and with Eq. 15 (that is, a given distance with a known force resistance, as in a race; see Eq. 17. With *P*_*i*_, *P*_*c*_ and *τ* individually obtained using RACLET. As described above, systematic and random errors, as well as the coefficient of correlation, root mean square error (RMSE) and comparison between the regression equation and identity line, were computed.

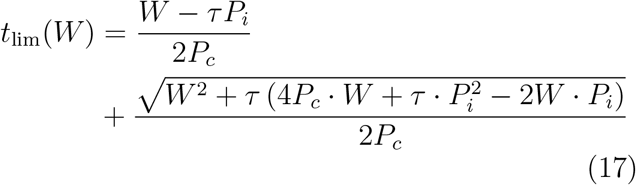

## Results

### Feasibility

The iso-cadence condition throughout the tests was verified, as the overall cadence was 80.6 ± 1.0 rpm (Fig. 6). The actual fraction of 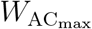 consumed during the RACLET was 57.7% *±* 19.7%. The highest RPE during the test was 16.6 *±* 1.7 and 12.0 *±* 1.6 at the end of the test.

The RACLET settings (10 sprints, sprint-time/total-time ratio = 10%, RACLET duration = 300 s,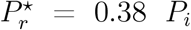) allowed an excellent goodness of fit for Eq. 4 from the maximum power data at 80 −rpm as a function of time (median *r*^2^ = 0.946, RMSE = 3.5%*P*_*i*_; A typical trace is displayed in Fig. 5A.).

### Reliability

The results of the reliability analysis are displayed in table 1. The RACLET parameters *P* ^⋆^ and *s* showed an excellent (ICC*>*0.985) associated with the relative consumption of 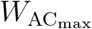 with high reliability (ICC = 0.86). The parameters *P*_*i*_ and *P*_*c*_ showed excellent relative (ICC = 0.993 and 0.985, respectively) and absolute (TEM = 0.9 and 4.2%, respectively) reliabilities. The correlational analysis of *P*_*c*_ between RACLET (A) and (B) is shown in Fig. 7A.. The model’s time constant exhibited weaker reliability (*τ* : ICC=0.698, TEM = 23.7% and 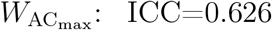 and TEM= 22.6%).

**Table 1.** Test-retest of the RACLET parameters and their reliability statistics.

| n=23 | RACLET (A) | RACLET (B) | CM | TEM | TEM% | ICC | r | p |
| --- | --- | --- | --- | --- | --- | --- | --- | --- |
| $P^*$ [W] | 424.94±96.59 | 421.00±94.24 | -3.95 [-5.59;-3.05] | 7.87 [6.09;11.14] | 1.9 | 0.993 [0.982;0.997] | 0.993 | <0.001 |
| $s$ [W·s<sup>-1</sup>] | 1.29±0.28 | 1.27±0.28 | -0.02 [-0.03;-0.01] | 0.03 [0.02;0.05] | 2.5 | 0.985 [0.963;0.994] | 0.987 | <0.001 |
| $P_i$ [W] | 1108.44±241.76 | 1108.98±237.54 | 0.55 [0.42;0.78] | 10.42 [8.06;14.75] | 0.9 | 0.998 [0.996;0.999] | 0.998 | <0.001 |
| $P_c$ [W] | 272.72±63.80 | 268.47±64.69 | -4.25 [-6.01;-3.29] | 11.48 [8.88;16.25] | 4.2 | 0.967 [0.925;0.986] | 0.968 | <0.001 |
| $\tau$ [s] | 19.91±5.82 | 21.69±10.77 | 1.78 [1.37;2.51] | 4.72 [3.65;6.67] | 23.7 | 0.698 [0.416;0.858] | 0.841 | <0.001 |
| $W_{AC_{max}}$ [kJ] | 16.15±4.02 | 17.61±7.53 | 1.47 [1.13;2.08] | 3.65 [2.82;5.17] | 22.6 | 0.626 [0.307;0.820] | 0.763 | <0.001 |
| %$W_{AC_{max}}$ | 57.62±18.56 | 57.87±20.88 | 0.25 [0.19;0.35] | 7.53 [5.82;10.66] | 13.1 | 0.860 [0.698;0.938] | 0.861 | <0.001 |

$P_i$ , $P_c$ , $\tau$ and $W_{AC_{max}}$ are the initial, critical power, time constant and impulse reserve (critical power model's parameters). $P^*$ , $s$ and % $W_{AC_{max}}$ are the starting power, power-time slope and final $W_{AC}$ consumption (RACLET's parameters). CM: Change in the Mean; TEM: Typical Error of Measurement in absolute unit or %; ICC: Intraclass Correlation Coefficient; r and p: Pearson's correlation coefficient between Test and Retest and the associated p-value.

**Figure 7.**
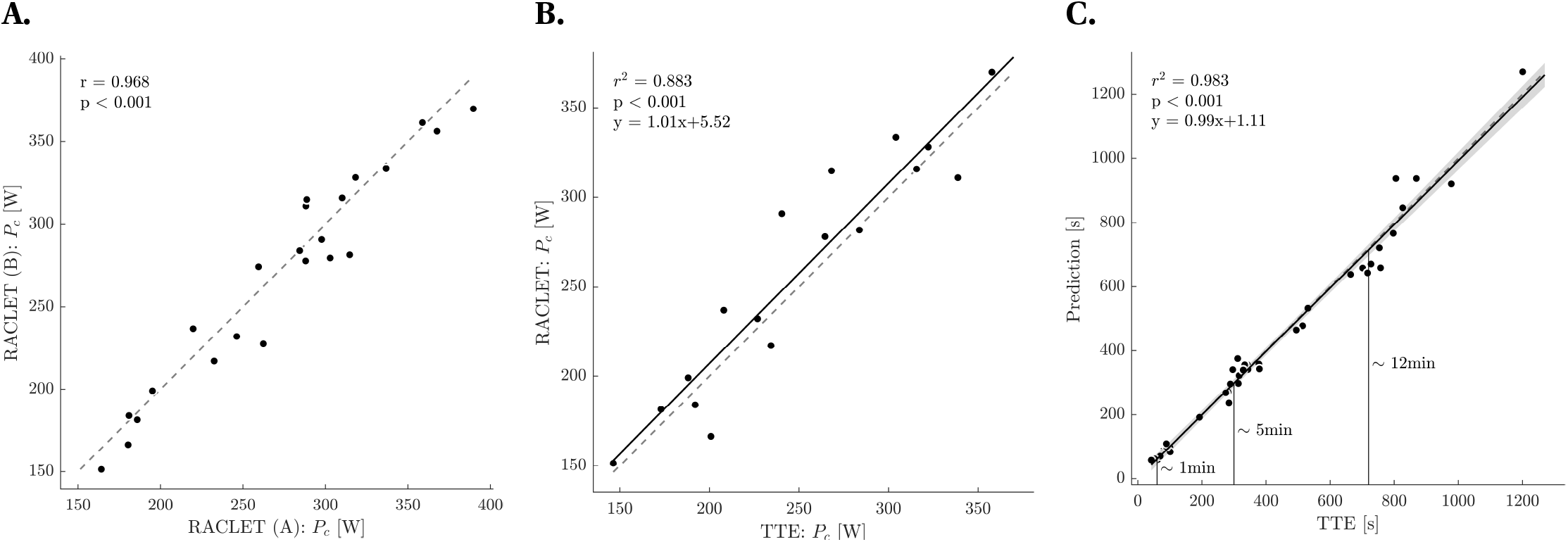
Correlation analysis for the critical power (A) Test-retest reliability using the RACLET protocol; (B) Validity comparison between RACLET and time-to-exhaustion (TTE) method; (C) Predictive model performance for TTE. Each dot represents a participant. Identity line (dashed), regression line (solid), and 95% confidence interval (shaded area) are shown.

### Validity

#### Concurrent agreement

The concurrent agreement between RACLEt and TTE parameters are shown in table 2. *Pi* and *P*_*c*_ showed good agreement with the gold standard method (systematic error: 1.7 and 3.0%; random error: 3.0 and 9.1%; Pearson’s correlation coefficient: 0.981 and 0.940; regression line not different from identity line). The correlation analysis of *P*_*c*_ between RACLET and TTE is shown in Fig. 7B.. In contrast, *τ* and 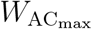 presented a random error of > 30% and a non-significant correlation between the RACLET and TTE.

**Table 2.**
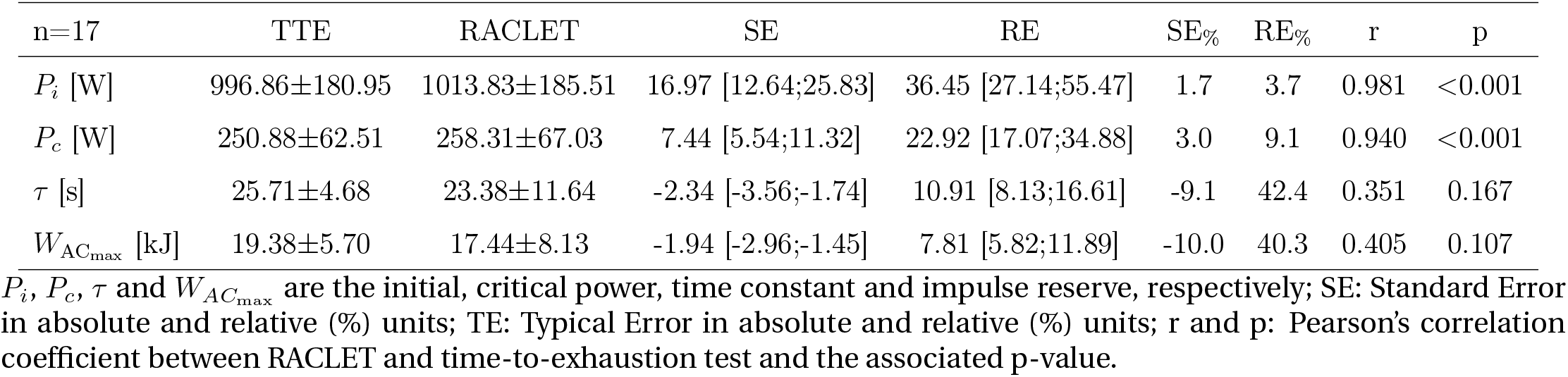
Validity statistics of the critical power’s parameters determined from the RACLET compared to the time-to-exhaustion tests.

#### Prediction

The time-to-exhaustion (*t*_lim_) measured during the experiment were very well predicted by the parameters obtained from the RACLET (Fig. 7C.). Actual and predicted durations presented an excellent correlation (r=0.992; p*<*0.001; RMSE=10.4%), with a systematic error of 2.13 s (−0.6%) and a random error of 39.3 s (10.3%) in table 3. When the relative prediction error was reported for each of the evaluated durations, the systematic errors were 5.5%, 0.6%, and −1.6%, and the random errors were 12.%, 9.2%, and 8.2% at 1, 5, and 12 min, respectively.

**Table 3.** Time-to-exhaustion prediction capacity statistics of the critical power model from the individually fitted *P*_*i*_, *P*_*c*_ and *τ*.

| n=17 | TTE | Prediction | SE | RE | SE% | RE% | RMSE | RMSE% | r |
| --- | --- | --- | --- | --- | --- | --- | --- | --- | --- |
| ~1 min [s] | 73.55±16.26 | 77.57±15.39 | 4.03 [2.95;6.35] | 8.85 [6.48;13.95] | 5.5 | 12.0 | 8.54 | 11.6 | 0.845 |
| ~5 min [s] | 310.93±46.00 | 312.87±51.93 | 1.94 [1.42;3.06] | 28.49 [20.86;44.93] | 0.6 | 9.2 | 29.45 | 9.5 | 0.837 |
| ~12 min [s] | 755.78±181.32 | 743.42±211.95 | -12.36 [-19.49;-9.05] | 61.66 [45.14;97.24] | -1.6 | 8.2 | 59.48 | 7.9 | 0.963 |
| All [s] | 380.08±304.96 | 377.96±304.90 | -2.13 [-2.69;-1.76] | 39.33 [32.56;49.68] | -0.6 | 10.3 | 39.70 | 10.4 | 0.992 |

SE: Standard Error in absolute and relative (%) units; RE: Random Error in absolute and relative (%) units; RMSE: root mean square error in absolute and relative (%) units; r: Pearson's correlation coefficient between predicted and observed time-to-exhaustion.

## Discussion

The purpose of this study was to design a new test, the Ramp Above Critical Level Endurance Test (RACLET), and assess its feasibility before investigating its interday reliability, concurrent agreement with the gold standard method and predictive capacity. The main results were as follows: (i) the participants reached a moderate level of fatigue during the RACLET (57.7% consumption of 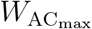; highest mean RPE of 16.6 ± 1.7); (ii) *P*_*i*_ and *P*_*c*_ showed excellent inter-day reliability (ICC*>*0.985) and an acceptable reliability was found for *τ* (ICC = 0.698); (iii) concurrent agreement was excellent to good for *P*_*i*_ and *P*_*c*_ (RE = 3.7 and 9.1%, respectively), but moderate for *τ* (RE = 42.4%); the RACLET capacity to predict time-to-exhaustion duration was excellent (RE = 10.4%).

First, we succeeded in configuring an iso-cadence (80.6 ± 1.1 rpm) RACLET such that it led to a moderate level of fatigue, as attested by the RPE being only 12 at the end of the test and the fraction of 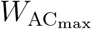 consumed of 57.7% in average. This proportion is close to that estimated *a priori* (50%) based solely on the literature data and Eq. 9. The minor difference could be due to a slightly different population from that used to calibrate the test (Dorel et al., 2010; Vanhatalo et al., 2007), but also to intermediate sprints which are not considered a prior in the estimation of 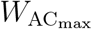 consumption, but which still contributed to fatigue. Nevertheless, the proposed method to set P⋆ and s seems sufficiently accurate to enable the RACLET parameters to be adapted to other populations with different physical characteristics.

Second, reliability is a required element to enable a confident use of test results. As expected and already widely reported, the initial maximum power capacity *P*_*i*_ proved to be reliable with extreme accuracy (ICC = 0.998; TEM = 0.9%). A key result of this study was that the critical power *P*_*c*_ also showed excellent reliability (ICC = 0.967; TEM = 4.2%; r = 0.968). These reliability indices are in line with those found in the literature for TTE and all-out methods (ICC range: 0.91-0.96; r range: 0.93-0.96; CV range: 3.0 - 5.4%; Byrd et al., 2021; Dekerle et al., 2014; Gaesser and Wilson, 1988; Nebelsick-Gullett et al., 1988). 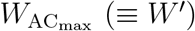 presented a lower reliability (ICC = 0.626; TEM = 22.6%; r = 0.763). This is slightly below the reliability values previously reported for historical methods (ICC range: 0.79-0.98; r range: 0.78-0.79; CV range: 5.4 - 18.6%; Byrd et al., 2021; Dekerle et al., 2014; Gaesser and Wilson, 1988; Lucas et al., 2014). The parameters *τ* and 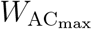 have an inter-individual variance of the order of magnitude of the random noise, which probably led to a moderate signal-to-noise ratio and could explain an important part of the low ICC observed. Nevertheless, RACLET appears to be a test that reliably determines the critical power model’s parameters. Moreover, improving the reliability of 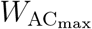 using other RACLET parameter settings (for instance, a higher fraction of 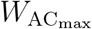 consumed) could be further explored.

When comparing the proposed RACLET to the TTE gold standard method, a very low systematic error and a low random error were observed for the initial and critical power (*P*_*i*_ SE_%_±RE_%_ = 1.7% ± 3.7%; *P*_*c*_ SE_%_±RE_%_ = 3.0%±9.1%). This typical error can be attributed, at least in part, to the reliability of each method and falls within the order of magnitude of typical errors previously reported for traditional method, such as the 3-min all-out test (2.8 - 7.76%; Clark et al., 2016; Wright et al., 2017). For *τ* and 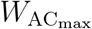 the systematic errors were moderate (−9.1 and −10.0%, respectively). The random error was significantly higher, ≈ 40%, for both. Again, given the lower reliability of these parameters (*vide supra*), this result is regularly reported when comparing other tests (e.g. 3min all-out, field testing) with the TTE method (for *W ′*, r range: 0.16 - 0.84 not always significant; random error range: 15.5 - 40.0%; Clark et al., 2016; Karsten et al., 2015; Wright et al., 2017). Therefore, the validity of RACLET appears to be very good for the power parameters *P*_*i*_ and *P*_*c*_, and moderate for *τ* and 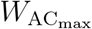, yet within the magnitudes observed for other widely used tests, such as the 3-min all-out test.

In addition to the reliability of parameter estimation and its agreement with reference methods, a crucial element in the use of a model is the validation of its predictive capacity. For the critical power model, the most frequent readout consists of predicting the duration of a constant power effort before reaching exhaustion or the time taken to cover a given distance or, on a stationary cyclo-ergometer, for a given work (i.e. time-trial; Muniz-Pumares et al., 2019). For each participant, we predicted the times of exhaustion for three different powers higher than *P*_*c*_ and their corresponding work production. All predictions were excellent (r=0.992; p*<*0.001; systematic error of 5.5%, 0.6%, −1.6% for the shorter, intermediate, and longer efforts, respectively (Fig. 7C.). These results are among the best predictions found in the cycling literature (*r* range: 0.87 - 0.99; CV range: 3 - 22%; Chidnok et al., 2013; Jones et al., 2008; Nicolò et al., 2017; Vanhatalo et al., 2011). Therefore, RACLET identifies a set of parameters *P*_*i*_, *P*_*c*_, and *τ* that allow the model to be applied with an excellent predictive capability.

The results of this study provide evidence that the RACLET can be used with confidence, as a feasible, reliable and valid test. Although the results could differ in other populations or under different experimental conditions, it is reasonable to think that RACLET can be used on a wide scale because of its fundamental rationale built on the definition of critical intensity (the heavy/severe intensity domain boundary). However, it is crucial to remember that the feasability/validity of this test is closely tied to the quality of the target power tracking, the maximum power measurements, and the setting of specific parameters of RACLET (*P* ^⋆^, slope *s*, test duration *t*_*R*_). Those intending to use this test should focus on these aspects. An improperly configured test, whether too easy or too difficult, a power developed far from the target or too noisy maximal capacities, will prevent from consistent, reliable and valid results or the test from being completed. Note that reliable and valid output data obtained here were associated with high goodness of fit of the model to maximal power over time (median *r*^2^ = 0.946), and a very good respect of the power target (reaching here via a good respect of iso-cadence condition against a decreasing friction, 80.6 *±* 1.0 rpm).

To assist anyone interested in implementing the RACLET, all the procedures for designing and analysing the test are detailed and available for free in Excel file and Matlab code on the FoVE research group’s Gitlab: https://gricad-gitlab.univ-grenoble-alpes.fr/fove/methods/raclet.

## Conclusion

### RACLET - Power

Based on a mathematical model previously developed by our team, this study proposes and validates a new single test: the Ramp Above Critical Level Endurance Test. The model’s parameters are the initial maximum power *P*_*i*_, the critical power *P*_*c*_ land a characteristic time *τ* (from which we can also determine 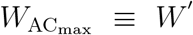), which proved to be reliable, valid, and as predictive as the gold standard methods. The major advantage of this method is that it does not lead to exhaustion (≈ 50% fatigue) and only requires maximal effort to be performed in very short intervals (< 3 s). This overcomes the major limitations of historical methods (multiple time-to-exhaustion or 3-min all-out test). It also opens the door to the evaluation of parameters of the critical power model that were previously very difficult, if not impossible, including the evaluation of vulnerable populations, frequent longitudinal monitoring of athletes, or in a research context, the study of parameters imposing a very large number of experimental conditions, (i.e. such as the evaluation of cadence conditions effect to access the critical power-velocity relationship).

## Abbreviations

*α*: *P*_*c*_ expressed relatively to *P*_*i*_
*β*: *W*_AC_ expressed relatively to 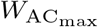
*γ*: *t*_*c*_ expressed relatively to *t*_*R*_
Γ: Torque
Γ_0_: Maximal theoretical torque at null cadence, without fatigue
Γ_*c*_: Critical theoretical force
Γ_*i*_: Initial maximal theoretical torque
Γ_max_: Maximal theoretical torque
Γ^⋆^: Torque at the start of RACLET
*ω*: Cadence
*ω*_0_: Maximal theoretical cadence at null torque, without fatigue
*P*: Power
*P*_*c*_: Critical theoretical power
*P*_*i*_: Initial maximal theoretical power
*P*_max_: Maximal theoretical power
*P*^⋆^: Power at the start of RACLET
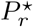 *P*^⋆^: expressed relatively to *P*_*i*_
RACLET: Ramp Above Critical Level Endurance Test
*S*: Torque ramp slope of RACLET
*s*: Power ramp slope of RACLET
*τ*: Typical time of fatigability
*t*_*c*_: time at which critical power is reached
*t*_lim_: Duration of Time to exhaustion
*t*_*R*_: total duration of RACLET
TTE: Time-To-Exhaustion
*W*: Work done during time-to-exhaustion
*W*_AC_: Work done Above Critical power
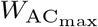 Maximal *W*_AC_: that can be done

## Acknowledgments

Support was provided by the French National Agency [ANR-22-CE14-0073] to B. Morel. The authors have no conflicts of interest relevant to the content of this article. The results of this study are presented clearly, honestly, and without fabrication, falsification, or inappropriate data manipulation.

